# Too Much Information Is No Information: How Machine Learning and Feature Selection Could Help in Understanding the Motor Control of Pointing

**DOI:** 10.1101/2022.10.08.511392

**Authors:** Elizabeth Thomas, Ferid Ben Ali, Arvind Tolambiya, Florian Chambellent, Jérémie Gaveau

## Abstract

The aim of this study was to develop the use of Machine Learning techniques as a means of multivariate analysis in studies of motor control. These studies generate a huge amount of data, the analysis of which continues to be largely univariate. We propose the use of machine learning classification and feature selection as a means of uncovering feature *combinations* that are altered between conditions. High dimensional electromyograms (EMG) vectors were generated as several arm and trunk muscles were recorded while subjects pointed at various angles above and below the gravity neutral horizontal plane. We used Linear Discriminant Analysis (LDA) to carry out binary classifications between the EMG vectors for pointing at a particular angle, versus pointing at the gravity neutral direction. Classification success provided a composite index of muscular adjustments for various task constraints – in this case, pointing angles. In order to find the combination of features that were significantly altered between task conditions, we conducted a post classification feature selection i.e. investigated which combination of features had allowed for the classification. Feature selection was done by comparing the representations of each category created by LDA for the classification. In other words computing the difference between the representations of each class. We propose that this approach will help with comparing high dimensional EMG patterns in two ways; i) quantifying the effects of the entire pattern rather than using single arbitrarily defined variables and ii) identifying the parts of the patterns that convey the most information regarding the investigated effects.

## 1.0 Introduction

Movement takes place through the contraction of several muscles which then causes the displacement of several body segments. Much pain and energy therefore goes into the simultaneous collection of the variables connected with motor control studies. Curiously despite the energy invested in the synchronization of the data collection and the big data tables created by such experiments, the analysis of it largely takes place in a univariate manner. In this study we propose the use of Machine Learning classification as a technique able to provide insights concerning global features of the voluminous datasets created from motor control studies. The use of these techniques with movement data is not new. However, it has been principally in the realm of application, largely unconcerned with questions concerning underlying mechanisms (Côté-Allard, 2019; Paranjauli et al, 2019; Phinyomark and Scheme 2018; Xiong et al, 2021; Labarrière et al, 2020). Compared to this, there are few studies using Machine Learning dedicated to the purpose of understanding the mechanisms underlying movement. There are many advantages to be gained from exploring this technique for understanding motor control. First, Machine Learning allows for the combination of variables in analyses hence providing the means for a global view. This is similar to the manner in which humans make decisions – weighing the input from a combination of features before arriving at a conclusion (Drugowitsch et al, 2014; Mercier and Cappe, 2020). Second, linear or nonlinear feature combinations of variables could allow for significant differences in cases where they would not individually. Thirdly, with a technique of classification and then feature analysis, we are using an approach that is more in keeping with the current spirit of ‘big data’. Rather than imposing previously held views concerning which variables are important, we allow this information to emerge from an understanding of which variable combination is important for classification. This process of identifying the features which are important for classification is an important field in Machine Learning called feature selection (Guyon and Elisseef, 2003; Saeys et al, 2007; Hira and Gillies, 2015; Jovic et al, 2015; Venkatesh and Anuradha, 2019).

The motor control task studied in this paper was one of pointing in different directions. Subjects pointed in the gravity neutral horizontal dimension of 90° and then at 180° (vertically downwards), 135°, 45° and 0° (vertically upwards) (figure 1). These four directions entailed gravity constraints of varying degrees. Since the electromyographic activity (EMG) of nine muscles were recorded at high frequency (1000Hz) during pointing, each movement was associated with several high dimensional vectors for each subject. Although they have much to offer in the way of understanding motor control, EMG signals are not used as often as kinematic data in part due to their high complexity and intra and inter subject variability (Latash et al, 2008; Hagen & Valero-Cuevas, 2017). The experimental results describing the kinematic aspects of this current study have been published (Gaveau et al, 2016).

**Figure 1:**
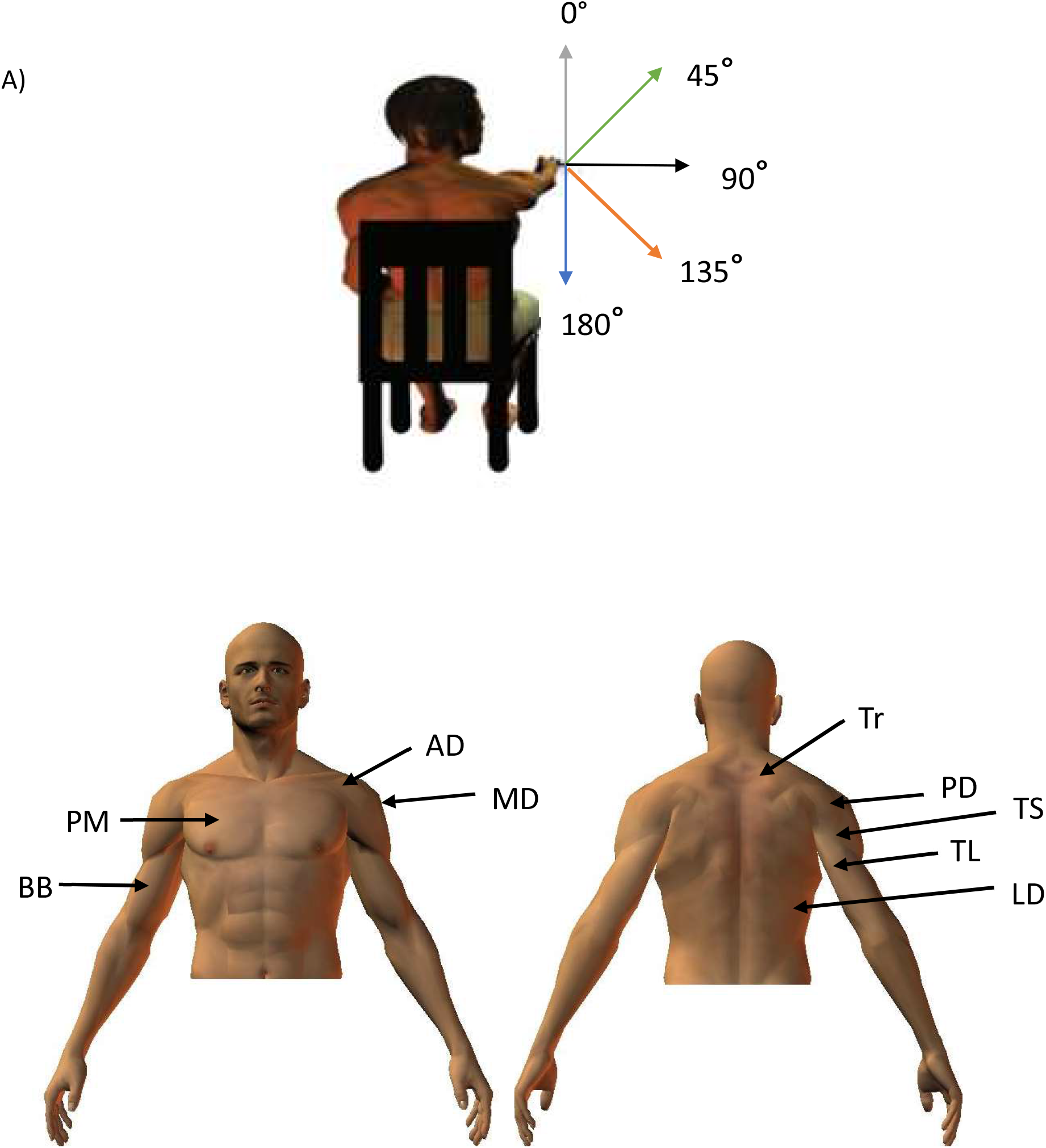
A) Participants were seated with a hand held horizontally in front of them. They were then asked to point to targets placed at 180°, 135°, 45°and 0°B) Approximate positions of recorded muscles.

Machine learning was used to analyse the data as it would provide us with a means of conducting a simultaneous analysis of all the recorded EMGs. The task was approached as a binary discrimination task with each case being the classification of EMGs in the gravity neutral direction of 90° compared to pointing at 180°, 135°, 45° or 0°. Importance was given to the use of Linear Discriminant Analysis (LDA) as the algorithm for classification. In LDA, the means and variance of each group are used to construct models of each group which are multivariate Gaussians. The probability of a data point belonging to one class or the other is then computed with the help of the Bayes factor (Grimm and Yarnold, 2006; Johnson and Wichern 2007; Izenman, 2013). This is a relatively old algorithm and several previous studies including some of our own have shown that more recent techniques like the kernel methods (Support Vector Machines belong to this group of methods) or random forest trees provide better classification (Han et al, 2017; Heung et al, 2016; Nair et al, 2010; Statnikov et al, 2008; Uddin et al, 2019). We nevertheless chose to work with this technique due to its ease of application especially with regards to feature selection. We also hope that its conceptual connection with classical univariate statistics would also encourage the greater use of machine learning in the basic science studies of motor control.

Once the classification has been performed we proceeded to identify the feature combination which had allowed for discrimination. This process of feature selection in the field of engineering is largely for the purpose of reducing the size of datasets and hence speeding up classification. These efforts therefore concentrate on picking the *minimum* number of features necessary for classification. In contrast, the application of such techniques to the basic sciences, would require an identification of several of the features/feature combinations which have been altered. Standard features selection techniques can be grouped into the categories of filter, wrapper and embedded techniques. Review articles which explain the central philosophies of each of these techniques can be found in several previous publications (Guyon and Elisseef, 2003; Saeys et al, 2007; Hira and Gillies, 2015; Jovic et al, 2015; Venkatesh and Anuradha, 2019, Remeseiro and Bolon-Canedo, 2019). Filter methods are usually pre machine learning methods which first assign a statistical score to each feature. They are generally used as a pre-processing step and selection of features is independent of any machine learning algorithms. The features are then ranked according to these scores and the features which are within a cutoff threshold are kept. These methods are often univariate. Some examples are the chi squared test and correlation coefficient score. Wrapper methods are based on the results obtained from machine learning algorithms. Through recursive processes, the method uses various search methods to assemble feature combinations which are then tested using the classification algorithm. Putting together the feature combinations can range from greedy algorithms like sequential forward selection or more optimised search methods like randomized hill climbing. In embedded techniques, the search for the optimal feature combination for classification is done as the machine learning algorithm is being constructed. Regularization methods are embedded feature selection methods. Such methods give a weight to each feature that the learning algorithm optimises, so that the unimportant features have a close to zero weight and are eliminated from the algorithm calculations. Examples of such algorithms are LASSO (Least Absolute Shrinkage and Selection Operator) (Tibshirani, 1996) and ridge regression (Hoerl and Kennard, 1970). The feature selection method used in this study would fall under the embedded method category. Classification begins with a full feature vector. The differences between the models of each class created by LDA is then used to find the group of important features. We chose this method because it was in keeping with intentions of finding feature combinations which are altered between task conditions and to avoid making pre-suppositions concerning altered features. A successful categorization indicates the presence of information which is discriminatory between two groups. The representation of each category can then be analysed for finding the most relevant features to classification.

There is a good deal of information available concerning EMG activation patterns during pointing in different directions. These studies have helped to tease out a portion of muscular control which is involved in postural aspects of the movement from the aspects which are specifically related to directional tuning. The former, tonic EMGs, do not scale with task constraints such as speed while the latter, phasic EMGs, do so. These studies have also identified muscular alterations for pointing direction (Buneo et al, 1994, 2008; Flanders et al, 1991, 1992, 1996). Work by the Gaveau and his collaborators has shown how differences in the kinematics and muscular activity of downward and upward pointing can be related to the minimization of effort (Gaveau et al, 2016, 2021; Poirrier et al, 2022a). The group also showed how this minimization is altered with age (Poirier et al, 2022b).

Almost all the studies in motor control are univariate, following the traditional pattern of picking variables based on pre-conceived notions and comparing them using traditional statistics. An important exception to this has been the work using principle component techniques and its variants. This technique has also provided a composite image of muscular alterations during pointing by finding components which capture most of the variance in the data. Data dimensionality reduction in this technique has focused on finding a few synergies that can then be scaled to reproduce EMG patterns for several constraints (d’Avella et al, 2010, 2011; Delis et al, 2018). The main difference between this sort of technique and the Machine Learning Classification and feature selection techniques, is the nature of the focus on the data. While the former is primarily concerned with correlations and global similarities, the latter is concerned with global differences. Here, to illustrate the novelty of the former technique, we use it to answer the following question. What are the muscular differences when pointing at a particular angle compared to pointing in a gravity neutral direction?

The aim of this study is to investigate how LDA classification can be used to pick out the muscle combinations and temporal segments of EMG data that are most pertinent to changing pointing direction. Since the technique is able to provide composite indices of performance such as classification accuracy, or distance between models, we also aim to use these as a guide on global trends in muscular modifications for changing pointing directions.

## 2. Method

The aim of this project is to use Machine Learning in order to understand how muscle organization changes as a function of pointing direction. In order to do so, we will investigate if the class representations created during Machine Learning can be exploited for finding the features/muscles that are adapted for pointing direction. To this end, we will:

- Describe how the kinematic and EMG data were collected. Markers placed on the arm recorded kinematic data and hence allowed the researchers to verify the direction of arm movement. This section will only be brief as a detailed description of the kinematic aspects of the current study have already been published (Gaveau et al, 2016).
- Describe the pre-processing of the EMG data
- Describe how the data was organized as input for the machine learning classification
- Provide a brief description of the LDA algorithm
- Describe the technique of feature selection through the comparison of models created of each class by LDA.

### 2.1 Data acquisition

The input dataset used for the current investigation came from a previous study (Gaveau et al, 2016) which had been approved by the regional ethics committee of Burgundy. Data collection was carried out in keeping with regional and international norms (Declaration of Helsinki, 1964). It consisted of kinematics values and EMG signals recorded from 11 right handed healthy participants of mean age 24 ± 3.2 years (9 female, 2 male) who had given their written consent for the study. Right handed preference was evaluated by the Edinburg test (individual scores >0.86; Oldfield 1971).

The participants pointed in different directions from a seated position. They were seated with a hand held horizontally in front of them. They were then asked to point to targets placed at 180°, 135°, 45°and 0°as shown in figure 1. The order of pointing in different directions was randomised. There were 9 trials for each angle. The movements of the arm were followed through the use of kinematic markers placed on the shoulders, arm and hand of the subjects. For our purpose, we were only interested in the hand marker placed on the first metacarpo-phalageal joint. The trajectory of the marker was recorded using an optoelectronic system (Smart BTS Italy) with 4 cameras sampling at 120Hz. The X, Y and Z coordinates of the markers throughout the pointing movement were stored to identify the direction of pointing. Movement onset and offset were defined as the time at which hand tangential velocity went above or fell below 5% of maximum hand velocity.

The EMG data was obtained from surface electrodes placed on 9 muscles using the Aurion, Zerowire EMG at a sampling frequency of 1000Hz. The muscles recorded from were the anterior deltoid (AD), posterior deltoid (PD), medial deltoid (MD), pectoralis major (PM), latissimus dorsi (LD), trapezius (Tr), biceps brachii (BB), long triceps (TL) and short triceps (TS). We applied the following standard procedures to the EMG data. They were band pass filtered with a bandwidth from 20 to 300Hz, using the ‘butter’ and ‘filtfilt’ functions in Matlab. Before integrating the signal using a sliding window of 5ms, each EMG signal was cut off 200ms before movement onset and at movement offset. Finally, EMG data were filtered one more time to obtain smooth patterns (low-pass 5Hz). Interpolation was used to place all the collected time series on a common time base. i.e., duration was normalised.

### 2.2 Classification

All the classifications carried out in this study were binary classifications. Muscular activity for pointing at 90 ° was chosen as a reference direction. This was done because horizontal arm movements are gravity neutral whereas movements in the other directions move the arm in or against the direction of gravity. So the binary classifications carried out were 90°vs 180°, 135°, 45° and 0°. A succinct description of the steps taken for the LDA analysis were the following

1. For each subject, normalize the EMG amplitudes for each muscle so that the maximum amplitude is 1 and the minimum is −1.
2. Link the EMGs for all the muscles of each trial to create input vectors
3. Divide the subjects into 5 folds
4. Keep 4 folds for training and the remainder for testing
5. Using the Matlab LDA algorithm, create representations of each class only using the 4 training folds.
6. Verify goodness of this representation by testing classification using the remaining test fold.
7. To obtain the LDA_*diff*_ vector subtract the representations of each category
8. Portions of the LDA_*diff*_ vector which have high a amplitude indicate importance in classification. A cutoff margin can therefore be progressively lowered to pick out the most important portions of the LDA_*diff*_ vector.
9. Repeat the process with the next training and testing fold.

More details of each step are provided in the sections below.

#### 2.2.1 Input data organisation

Input data was organized as was done in many of our previous studies (Tolambiya et al, 2011, 2012; Nair et al, 2010; Laroche et al, 2014). The input vectors for classification algorithm were created through the concatenation of the kinematic or EMG time series for each trial. As explained in the data acquisition section above, the time series for each variable was a vector of 1000 elements. So for example, taking the case of the kinematic data, the input vector for each trial was 3000 elements long as we concatenated the X, Y and Z time series for a trial. Since a subject pointed to each direction 9 times, the total kinematic matrix for each subject was 9×3000 for one angle. The same was done for the EMGs. Since there were 9 muscles, the EMG vector for each trial was composed of 9000 elements and the EMG matrix for one direction was 9×9000 for each subject.

It is to be noted that throughout the study we kept the entire time series of any sensor rather than using condensed features such as averages or frequencies. We also did not use features extraction techniques like PCAs to reduce dimensionality. This was done so as to facilitate the task of identifying the features which are important for the classification, not only in terms of muscles but also in terms of *when* the important muscular modifications took place.

The study was done using 5 fold cross validation. Due to the awkward number of participants, four of the folds had 2 subjects while the final fold had 3. In accordance with cross validation protocol, every fold had some trials in which it was used as training data and others in which it was used in the test set. However, there were no trials in which data from one participant played both training and testing roles at the same time.

Accuracy was computed as the mean accuracy over all test sets. As is the case for most studies on Machine Learning the data was normalized so as to give equal importance to each muscle. For each individual and each muscle, the maximum EMG amplitude was given the value 1 and a value of −1 was assigned to the minimum. The normalization was done over all trials and separately for each class of binary classification. Note that this form of normalization allows each muscle to have equal importance in classification. It also preserves differences in EMG amplitude when pointing in different directions.

#### 2.2.2 Linear Discriminant Analysis for Classification and Feature Selection

Classification and feature selection was done using Linear Discriminant Analysis (LDA) (Grimm and Yarnold, 2006; Johnson and Wichern 2007; Izenman, 2013). If we have a p dimensional vector x, the class k to which it belongs can be determined by computing the posterior probability that it belongs to a class Y=k by using Bayes rule.

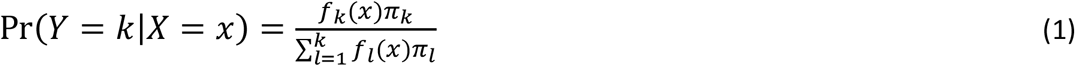

The variable *π_k_* is the prior probability for class k. It is computed as the proportion of samples belonging to class k

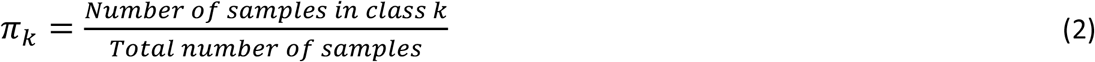

x would belong to the class Y=k with the highest posterior probability

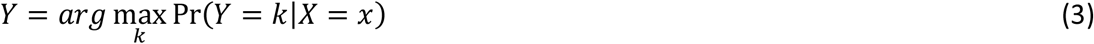

We can think of the function *f_k_*(*x*) as the model that LDA is using to represent each class *k*. It is the multivariate Gaussian of the data in each class

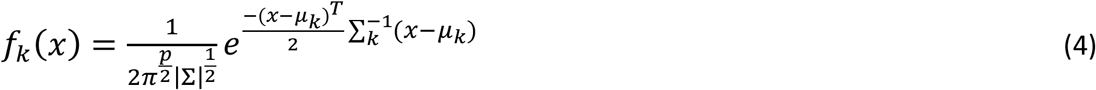

Where Σ is the *p* x *p* covariance matrix of X.

Feature selection was done by computing the variable *LDA_diff_*, obtained by comparing the models of each class created by LDA. This was the difference between the mean vector of each category. So if we were comparing two task constraints *k1 and k2*

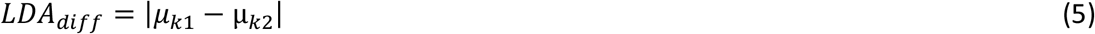

The cutoff threshold for finding the *High Diff* and *Low Diff* vectors from *LDA_diff_* was found in an iterative manner. It started at a high threshold value and was lowered step by step until the classification using the features above threshold was higher than what was obtained using the features below. Vectors that contain features above the cutoff threshold were called *High Diff* vectors, and those below, *Low Diff* vectors. Muscles which contained parts of the *High Diff* or *Low Diff* vectors were called *High Diff* or *Low Diff muscles*. The capacity of these EMG vectors to predict task constraints is reported in the Results section.

To obtain an idea of the composite difference between classes, we also computed another variable which we called the *LDA_distance_*. It was the sum of all the values in *LDA_diff_*.

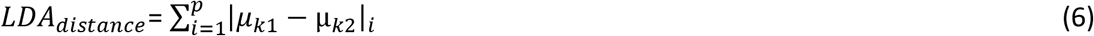

The LDA algorithm was implemented using Matlab (The MathWorks, Inc., Natick, Massachusetts, United States). Much care has to be taken in noting that there is no dimensionality reduction in this implementation of LDA. It is precisely this lack of dimensionalit’ redi ction which allows for feature selection by the simple step of subtracting the representations of each class.

#### 2.2.3 Statistics

In keeping with the categorical nature of classification, the statistics for the study were primarily done using the χ^2^ test i.e. By comparing the number of wrong versus right answers. When comparing the classification results for two different angles, we constructed contingency tables for the number of right versus wrong answers for each angle (Howell, 1992; Hinton, 1995). In the case of multiple comparisons, where we compared the classification accuracies at several angles, we applied the Bonferroni correction (Dunn, 1961; Goeman & Solari, 2014). Since this involve 4 pointing directions, results were significant if p<0.0125. As Figure 3B did not involve accuracies but a continuous variable, we used the Friedmann test followed by a planned comparison of the *LDA_distance_* for the classification at 180° versus 0°. The result was taken to be significant if p<0.05.

**Figure 2 :**
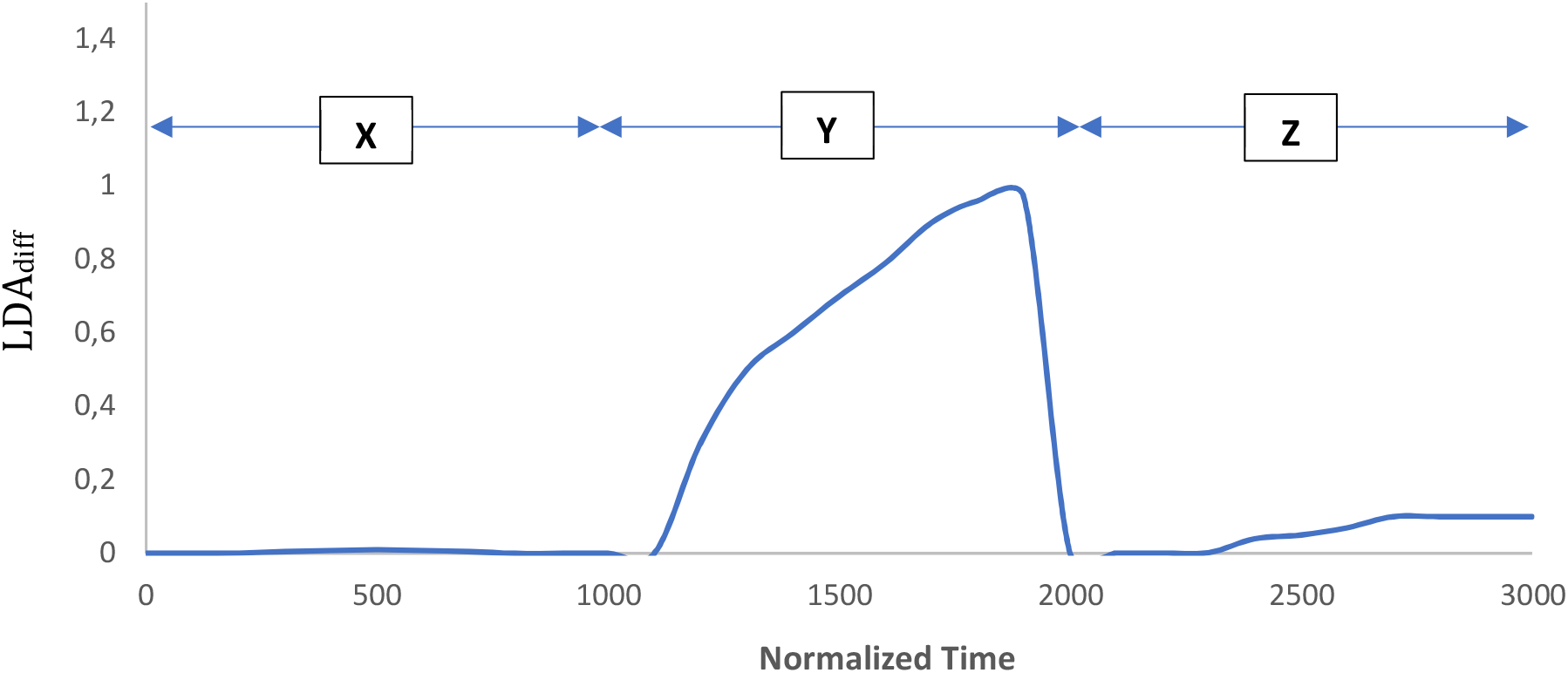
An example of a difference vector obtained by subtracting representations of each category following binary clasification of XYZ coordinates from kinematic markers from pointing towards 180° and 0°.

**Figure 3 :**
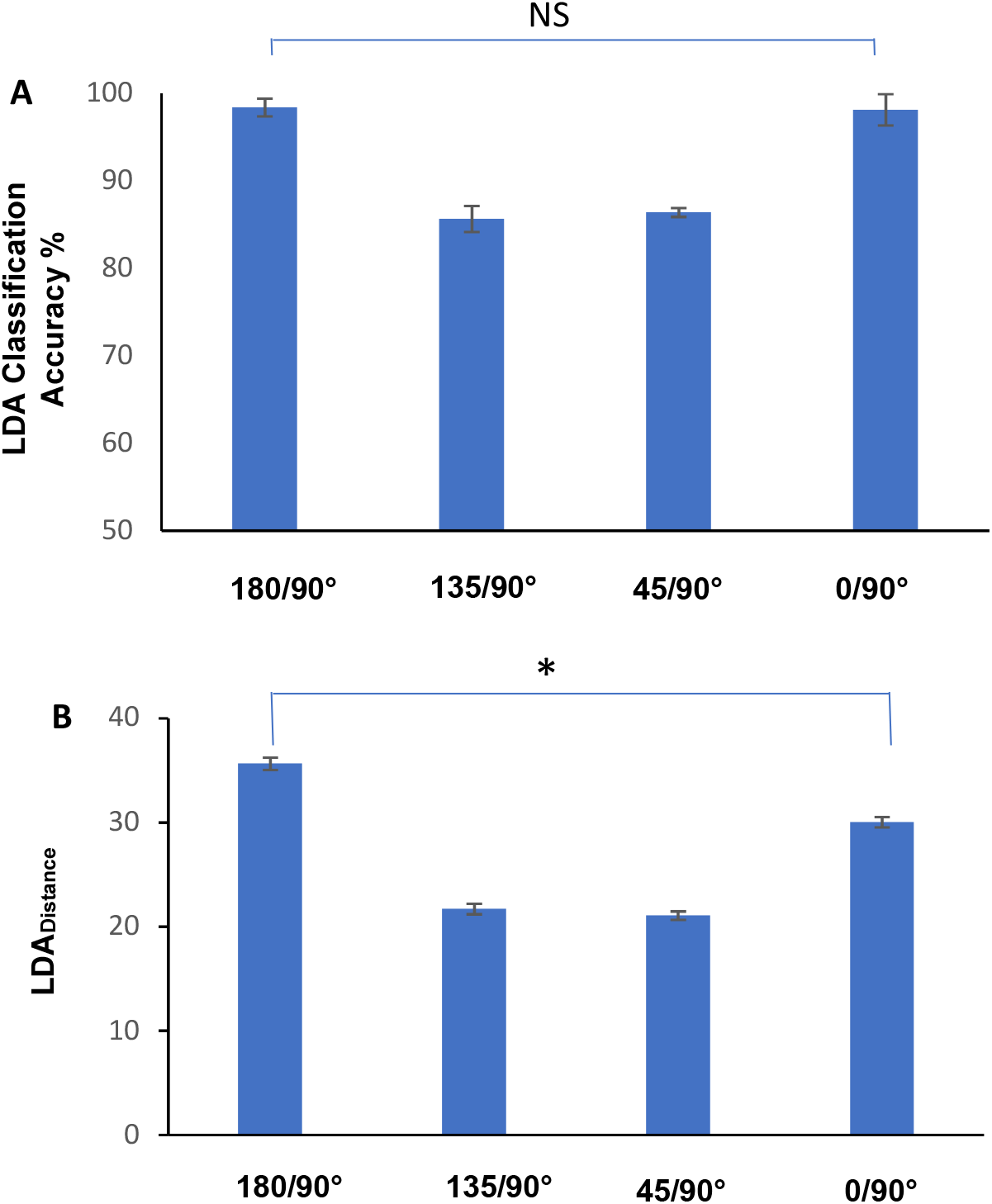
A) Classification accuracies from binary classifications between EMGs from pointing at 90° versus 180°, 135°, 45° and 0°. Of note was the lack of significant differences between pointing at 180° and 0° (p>0.05, χ^2^). B) The values of *LDA_distance_* obtained from the same classifications which were performed for figure 3A. While the classifications accuracies are similar for downward and upward pointing directions, the *LDA_distance_* variable shows that when compared to pointing in the gravity free horizontal dimension, EMG adjustments were greater when pointing downwards at 180° than when pointing upwards at 0° (p<0,05, Friedmann, followed by Wilcoxon planned comparison). All values represented in the figure are the means and ± standard error of the mean)

## 3. Results

In this section we will present the results from analysing the kinematics and EMG activities for different pointing directions. In each case, we will start out with the classification results of a full vector with all the features before comparing the models of each category to pick out the combination of features that are the most different between two task conditions, in other words, computing the *LDA_diff_* vector. This will then be followed by a comparison of the classification with the *High Diff* time series compared to classification with the *Low Diff* time series, the time series being either a muscle EMG or a kinematic time series. The investigation of this technique will first start with the easier example of automatic classification using the kinematic features from pointing in two clearly different directions. We will first predict using all the data from the kinematic markers if subjects had pointed upwards or downwards. Once we obtain success with this easier example which would serve as an essential proof of concept, we will move on to the more difficult task of automatic identification with the EMG time series for the pointing directions of 180°, 135°, 45° and 0°.

### 3.1 The kinematics of pointing upwards versus downwards

In this section we used the kinematic information from a marker placed on the index finger of subjects to classify if the subjects had pointed downwards (180°) or upwards (0°). The input vector for classification was a time series of the XYZ coordinates of this marker from the start of pointing movement to the maximum pointing amplitude. The classification accuracy obtained using LDA was 100%.

The next step was to use *LDA_diff_* to identify the features which were most relevant to this classification. Figures 2 displays the difference vector obtained from comparing the LDA created representations at 180° versus 0°. A visual inspection of the LDA difference vector showed a much bigger amplitude of *LDA_diff_* for the Y coordinate, while the differences for the X and Z dimensions were very low. According to our hypothesis, this indicated that the features most altered between the 2 pointing directions were those associated with the Y coordinate while the time series from the X and Z coordinates stayed relatively unchanged. This hypothesis was confirmed by an attempt at classification with the *High Diff Kinematic time series* (time series of Y coordinates) versus the *Low Diff kinematic time series* (time series of X and Z coordinates). Table 1 displays the classification results obtained from using the *High Diff* versus *Low Diff kinematic times series*. The results show that the classification accuracy and hence separation between the *High Diff kinematic time series* is higher compared to those for the *Low Diff kinematic time series*. This difference in separation was confirmed using the Support Vector Machine (SVM) and Learning Vector Quantization (LVQ) algorithms for classification. The difference in accuracy for these two kinematic series was found to be significant using all three Machine Learning algorithms (p<0.01, χ^2^ test) (Table 1).

**Table 1 :**
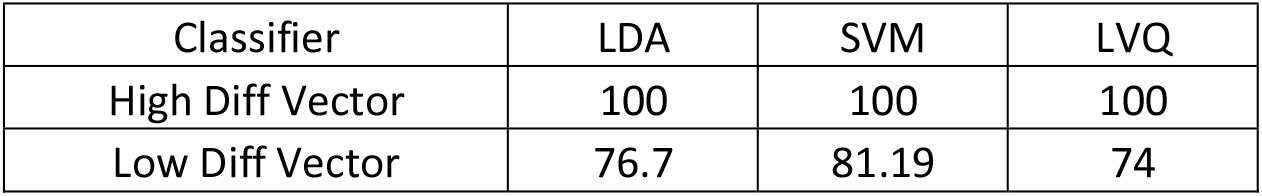
Comparison of classification accuracies using the *High Diff* versus *Low Diff kinematic time series*. All values in the table are mean percentages.

| Classifier | LDA | SVM | LVQ |
| --- | --- | --- | --- |
| High Diff Vector | 100 | 100 | 100 |
| Low Diff Vector | 76.7 | 81.19 | 74 |

This answer of which kinematic variable was most altered, was obviously correct as the only axis of movement was upwards and downwards. We only used it as an example and proof that examining *LDA_diff_* was able to reveal the most saliently altered variable in the time series.

### 3.2 Muscular alterations for pointing in different directions

Once we had tested our method of feature selection on the simpler kinematic case described above, we embarked on the more complex example of EMG activities for pointing in different directions. The directions tested were 180°, 135°, 45° and 0°. In each case, we conducted a binary classification in order to see which muscle combinations were significantly altered with respects to pointing horizontally at 90°, the gravity neutral direction. The organization of the input vectors and the sampling methods used for the classification are described in the Methods section above. The classification accuracies obtained using LDA can be seen in figure 3A. Following the classification, we once again computed the *LDA_diff_* to obtain an idea of the muscle combination which had contributed significantly to the classification and the moments at which they did so. Unlike the case with the kinematic features where one feature stood out in a very prominent manner, the situation was more complex in the case of the EMGs. In figure 4 we can see the difference vectors obtained from subtracting the representations for each binary classification. In the case of figures 4a and b, for downward pointing, it can be seen that the muscles that contributed the most to the difference vectors are the deltoid muscles and the trapezius. The *LDA_diff_* vector predicting pointing at 0° however (Figure 4d), would seem to indicate the involvement of a different muscle combination which had been modified with respects to pointing at 90°. This combination was more distributed, involving the Anterior and Medial deltoids, the Pectoralis major, Latissimus dorsi, Trapezius and Biceps brachii.

**Figure 4 :**
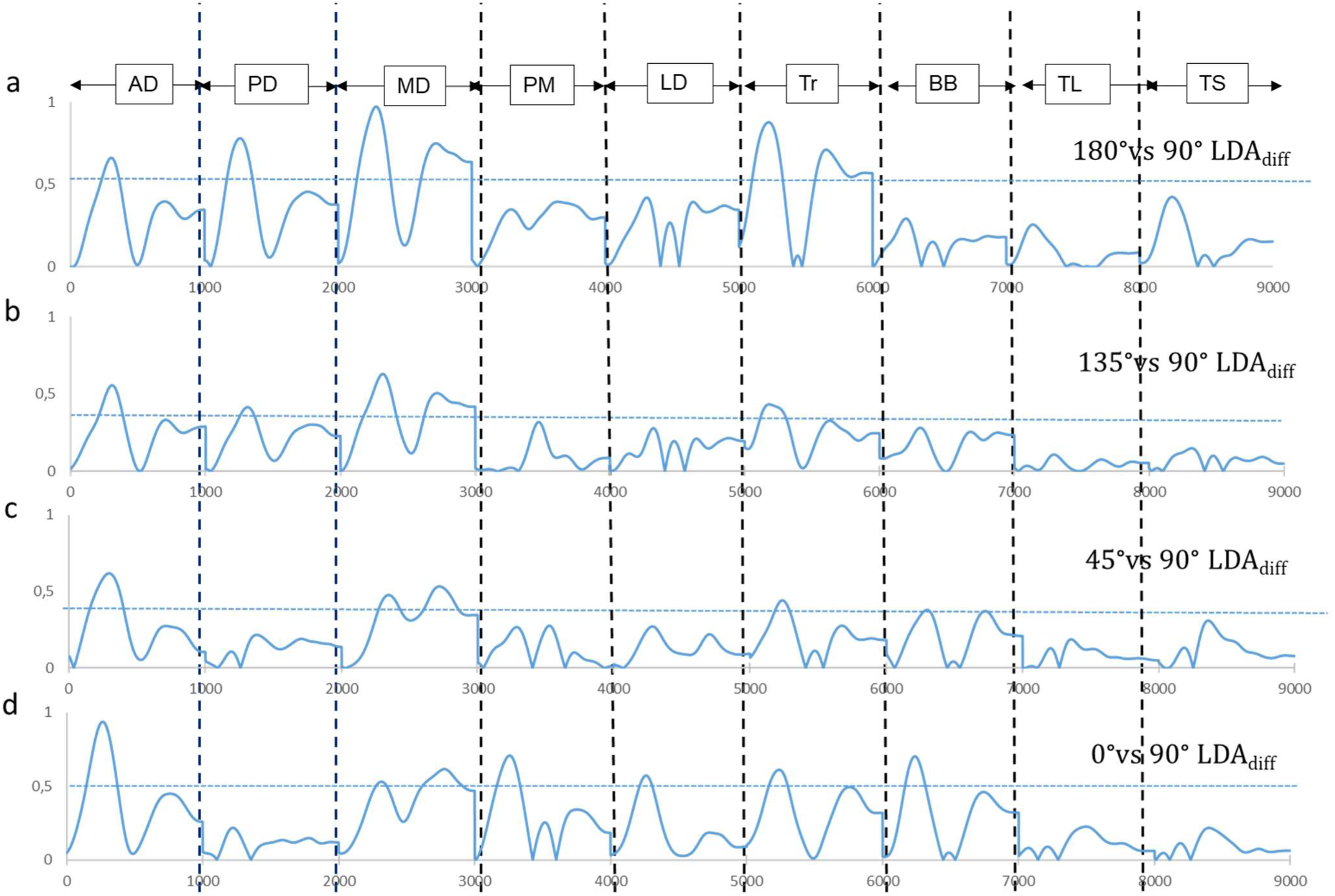
Difference vectors *LDA_diff_* obtained from the binary classification of input muscle vectors pointing horizontally versus at a) 180° b) 135° c) 45° d) 0°.

The next step taken was to apply the recurrent method described above to test if the combination of muscles with contributions from the *High Diff* vectors gave better classification results than those contributing from the *Low Diff* vectors. Table 2 displays the results of this test. The LDA classification with the *High Diff muscles* systematically gave better results than the *Low Diff muscles* for the binary classifications of pointing at 90° versus 180°, 135°, 45° and 0° (Table 2A, p<0.01, χ^2^ test).

**Table 2 :**
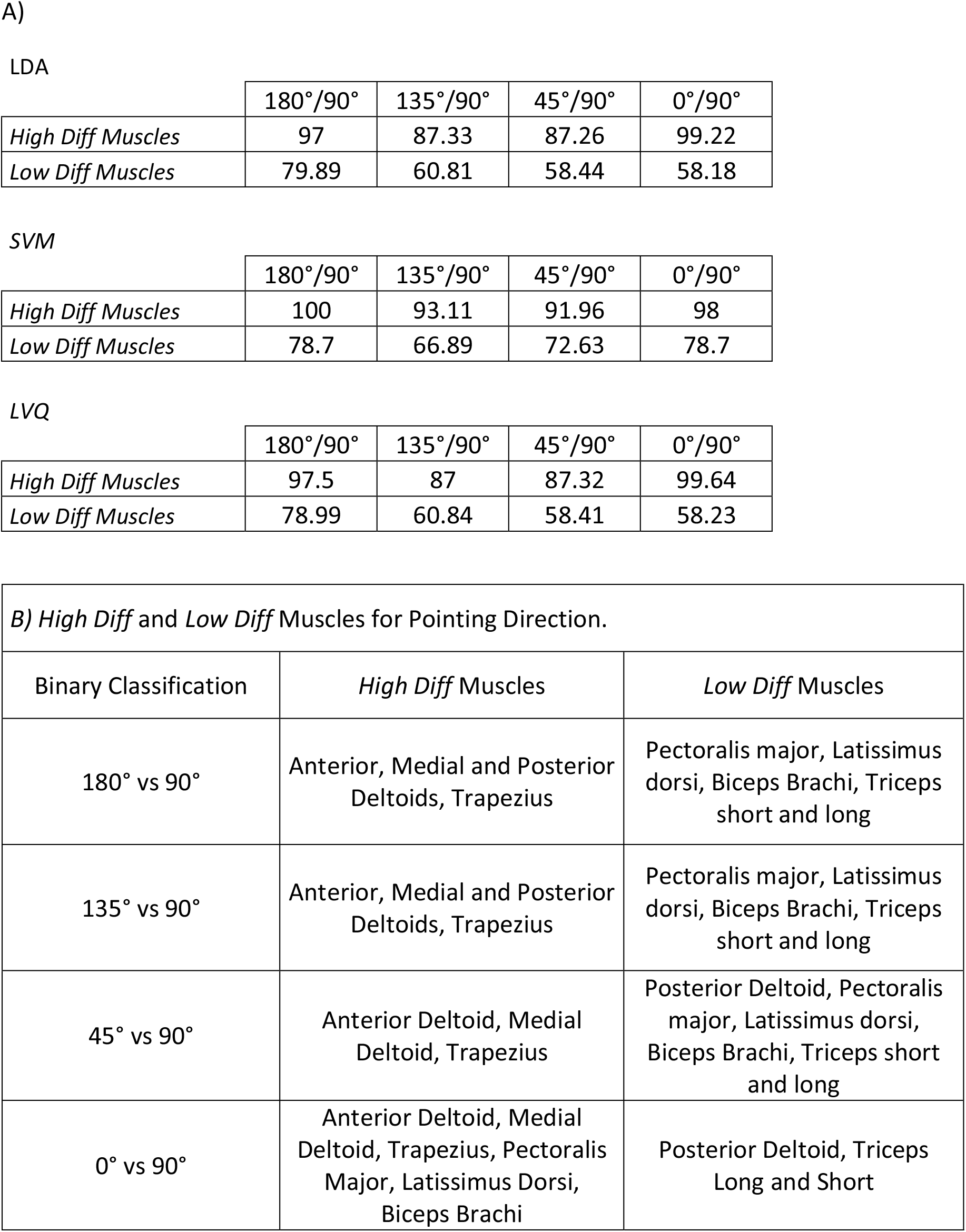
A) Comparison of classification accuracies using the EMGs from the *High Diff* versus *Low Diff muscles*. All values in the table are mean percentages B) Muscles identified as *High Diff* or *Low Diff* muscles for each pointing angle.

Next we tested whether the information obtained concerning the important components of the input vector from the model comparison with LDA, would hold for another classification technique. For this, we now compared classification accuracies between the *High Diff* and *Low Diff muscles* using the Support Vector Machine (SVM) and Learning Vector Quantization (LVQ) algorithm. Once again, as in the case of LDA, classification accuracies were significantly higher using the *High Diff* muscles indicating that this combination of muscles were more significantly altered for pointing in different directions (p<0.01, χ^2^ test).

In table 2B we have listed the *High Diff* and *Low Diff muscles* for pointing in various directions. They show that in all cases, the shoulder deltoid muscles were tuned to directional constraints. However, in comparison to the anterior and medial deltoid, the posterior deltoid did not play an important role in tuning towards the upward angles of 45° and 0°. Like the anterior and medial deltoid muscle, the trapezius muscle was important for modifications in all directions. In comparison to the other 3 angles, many more muscles were involved in the *High Diff* category for upward movement at 0°. Two additional trunk muscles, the latissimus dorsi and the pectoralis major played a more important role in this adjustment. For this direction, we also observed a bigger role for the biceps brachii.

### 3.3 The LDA Distance – A composite index of data separation

Classification accuracies provide a measure of data separability. High classification accuracies indicate that the data is highly separable and that the intra class variability is sufficiently low. On the other hand, once a certain amount of separability is present, classification accuracies would continue being high with a ceiling effect. In other words, there is more information to be gleaned from the *LDA_distance_* variable than from classification accuracies. In figure 3A, the classification accuracies at 180° and 0° were not found to be significantly different (p>0.05, χ^2^). This is not surprising as both accuracy values were similarly close to 100% (see Table 2). More precise information concerning data separation was obtained using the *LDA_distance_* variable. Figure 3B displays the mean values of the *LDA_distance_* obtained from the binary classifications of the EMGs from pointing at 90° versus 180°, 135°, 45° and 0°. While the classifications accuracies are similar for downward (180°) and upward (0°) pointing directions (figure 3A), the *LDA_distance_* values show that when compared to pointing in the gravity free horizontal dimension, EMG adjustments with respects to horizontal pointing were greater when pointing downwards at 180 degrees than when pointing upwards at 0 degrees (p<0.05, Friedmann, followed by Wilcoxon planned comparison).

### 3.4 The *LDA_diff_* vector and Temporal Features

Rather than using concise representations of muscle EMGs such as the means or the maximum amplitude, we chose to keep the entire time series of EMG activity. This was so that the LDA difference vector *LDA_diff_* would provide us with an understanding of the aspects of EMG activity which were altered not only in terms of amplitude but also in time. An inspection of *LDA_diff_* for all the cases of binary classifications performed in the study showed that the biggest values of *LDA_diff_* occurred in the first half of the movement. Once again, to test the idea that *LDA_diff_* provides an index of feature importance, we took one muscle, the anterior deltoid, and compared classification accuracies for all the binary classifications with the first half of *LDA_diff_ (First Half temporal vector*) compared to the second half (*Second Half temporal vector*). The results of these classifications can be seen in Figure 5. The mean classification accuracies for all pointing directions were found to be greater in the first half of pointing. These differences however were not found to be significant when pointing in the downward directions (135° and 180°). The story however was different when pointing in the upward direction (45° and 0°), hence indicating that the most discriminable adjustments for pointing in these directions was at the start of pointing rather than in the latter half (p<0.01,χ^2^ and Bonferroni correction).

**Figure 5:**
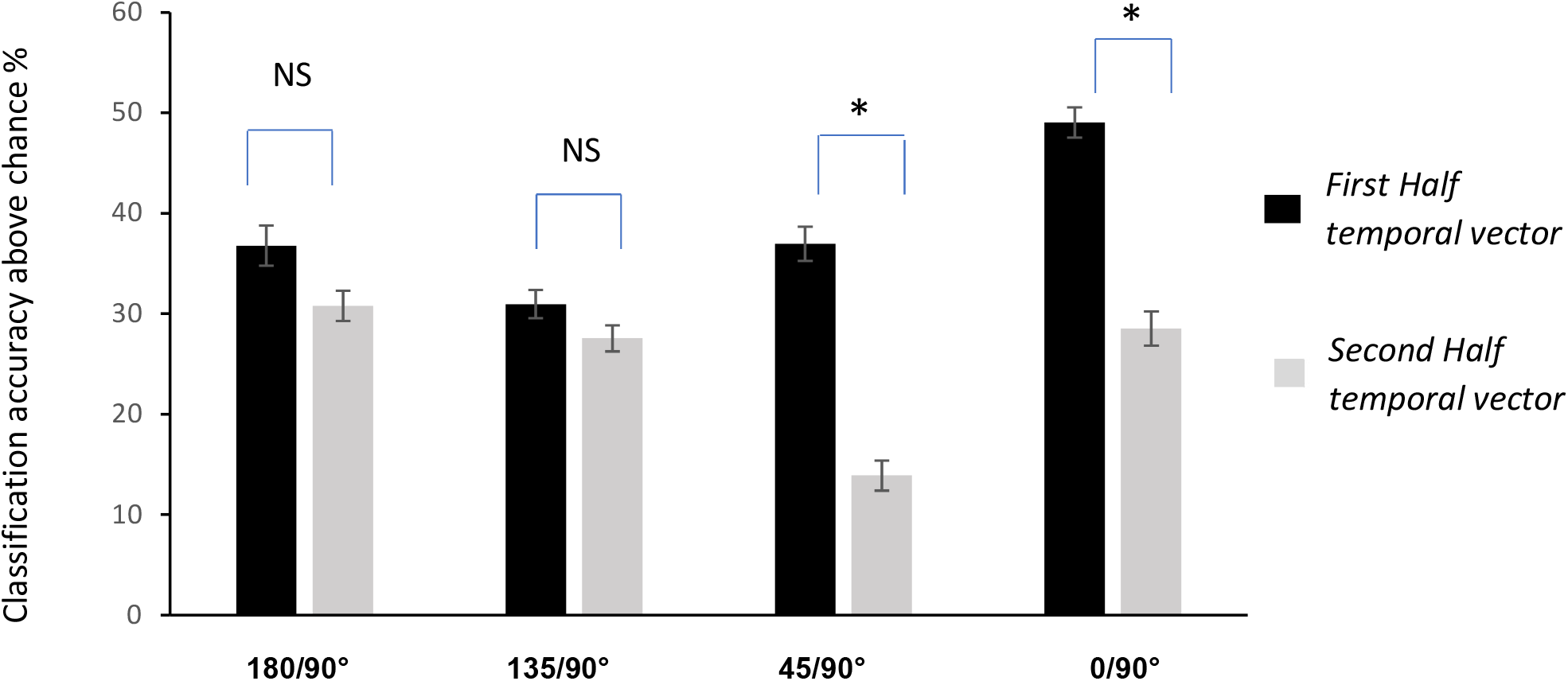
Classification accuracies obtained from the anterior deltoid EMG vector during the first and second half of the pointing movement. The figure shows that when pointing upwards, mean classification accuracies were higher in the first half of pointing. However, only the differences at 45° and 0° classifications were found to be significant (p<0.01,χ^2^ and bonferroni correction). Values displayed are mean ± standard error of the mean. Significant results are indicated with an asterisk, while NS indicates non-significant results.

### 3.5 Discussion

In this study, we have investigated the use of Machine Learning as an analytical tool which is appropriate for investigating the big amounts of data gathered in motor studies. Whether it be with experimental equipment in labs or phone applications which accompany people in their daily activities, advances in data collection has led to the creation of vast banks of information. The collection of this data is in keeping with a willingness to abandon an approach in which investigation is only carried out on a narrow set of pre-decided variables. This then opens up a new problem – among the big set of collected variables, which ones are important to the task at hand? We propose here, the use of Machine Learning classification and feature selection as a means of identifying this subset. In the sections below, we will start out by explaining how the focus of this paper is fundamentally different from many previous papers combining EMGs and Machine Learning where the emphasis was on application. We will go on to explain our choice of LDA as a Machine Learning algorithm. We will further discuss our choice of feature selection methods and compare it to previous techniques and finally, we will discuss the results from the project in the context of our current understanding of muscular contributions to pointing.

When it comes to motor activity, the primary use of Machine Learning has been in the field of engineering, many of these for the control of prosthetic limbs or for improvements in patient identification. So for example, Côté-Allard et al (2019) reported on how deep learning could be succesfuly used to recognize hand gestures. On a more challenging level, Parajuli et al (2019) wrote a review in which they described several studies that used Machine Learning to control hand prostheses in real time. Still within the framework of application, several articles on methodological issues with respects to the use of EMGs in Machine learning applications have been written by Phinyomark and Scheme (2018) as well as by Xiong et al, 2021). Just as in the case of upper limb prosthetics, the use of Machine Learning with EMG signals from lower limbs have contributed to the control of prosthetics. This was described in a review article by Labarrière et al (2020). A clinical study on how patients with arthritis could be identified through the use of lower limb EMGs and Machine Learning has been described by Nair et al (2010). Cheron et al (1996) described how lower limb kinematics could be mapped from lower limb EMGs using a recurrent neural network.

While the use of feature combinations in Machine Learning has proved to be useful in engineering applications, the approach can also be a way to tackle big data sets in basic research. By elucidating which features are important to discriminating task conditions, the technique can be used to identify the EMG or kinematic features which are the most altered between conditions. To our knowledge there has not been much research on the use of this approach in motor control. Some exceptions to this are studies by Tolambiya et al (2011) where the SVM was used to analyze EMGs during Whole Body Pointing. The study showed that the *combination* of postural rather than focal muscles provided a better prediction of pointing constraints, hence demonstrating that postural, rather than focal muscles, underwent greater modifications for several different variants of the Whole Body Pointing task. Using the same approach, Tolambiya et al (2012) showed in the anticipatory phase, that the *combination* of flexor rather than extensor muscles provided a higher than chance prediction between which of 4 different types of Whole Body Pointing tasks was about to be executed. This is indicative of differences in motor planning before the start of movement. In gait, Laroche et al (2014) showed that the thigh sagittal angle was able to provide a discrimination of patients with hip osteoarthritis versus control subjects as high as that of the combination of all other kinematic angles, hence indicating a high degree of modification in this angle for patients. These studies were able to exploit Machine Learning in the service of understanding how muscle and joint combinations contribute to motor control, hence presenting a departure from univariate studies. Nevertheless, the aforementioned studies on Machine Learning and EMGs are different from the current study in two important aspects. Firstly, the data combinations to be tested were decided upon *a priori* as opposed to this study in which the importance of variable combinations emerged from the feature selection that was done after (a *posteriori*) Machine Learning. Secondly, since classification in the previous studies was done with pre-selected groups, an ensemble view that allowed us to obtain an idea of the relative contributions of each muscle and phase with respects to the entire dataset as in Figure 4 was not obtained.

In this study we put an emphasis on the use of LDA as a classification algorithm. Several previous studies, including some of our own, have shown that other methods such as the kernel methods which include SVMs or random forest are more efficient classifiers (Díaz-Uriarte and Alvarez de Andrés 2006, Han et al, 2017; Heung et al, 2016; Nair et al, 2010; Statnikov et al, 2008; Uddin et al, 2019, Chen et al, 2020). Another example more pertinent to motor control is a study by Aeles et al (2020) in which they used the SVM to uniquely distinguish between 78 individuals using their EMG signatures (Aeles et al, 2020). Indeed table 1 and 2 of this study show that the SVM provides better classification. However such efficient classification would yield poor information regarding the proximity of EMG data for pointing in different directions as the SVM is capable of exploiting very small differences to yield high classification ie we would face a ceiling effect. Our choice was therefore to move ahead with LDA with which classification accuracy would provide more information concerning data overlap and because of the ease with which we were able to perform feature selection. This is also more in keeping with the spirit of explainable artificial intelligence (XAI) (Murdoch et al, 2019; Arrieta et al, 2020). Although it goes under the guise of this new name, XAI, the attempt to give a priority to understanding the factors which permitted the black box of artificial intelligence to achieve its classification and hence obtain an understanding of underlying mechanisms is not new (Chan et al, 2002, Nair et al, 2010; Tolambiya et al, 2011, 2012). It should be pointed out that even a classification of 80% yields a statistical significance of p<0.01 (χ^2^ test). The LDA algorithm, being a method which relies on the creation of multivariate Gaussians models of each class, allows for the subtraction of representations and a quick visualization of the role of each feature in distcrimination. This difference also provides a means of quantifying the distance between classes. As opposed to this, the SVM and Random Forest are methods that rely on a more distributed representation whereby this technique of representation subtraction cannot be used so readily. We have cited here several studies in which methods which are based on decision trees like Random Forest, achieve a superior performance. Once again, this method is not ideal when we are dealing with long time series (the concatenated EMGs) from which we do not wish to pick out particular features *a priori*.

While we were able to apply LDA to this problem of motor control with healthy subject in a repeated measures or paired protocol (same subjects pointing at all angles), it should be noted that not all data sets would be so linearly discriminable. This might especially be the case for studies with independent groups like patients and controls. Patient data also tends to have high variability. In such cases, if we are to follow the logic of post-classification feature selection, the characteristics of the separating surface can be used to select the most discriminating features (Guyon et al, 2002, Weston et al, 2003).

LDA has been used for feature selection using all three paradigms described in the Introduction section. An example of the filter method was an investigation by Lei et al (2012) where the Fisher criterion was used to select the most discriminative features before application of LDA for face discrimination. An example of the wrapper method applied with LDA can be seen in the study by Gayathri and Sumathi (2016) where feature combinations were assembled *a priori* and then tested with LDA. We did not apply either of these techniques as they were not in keeping with our wish to have a method in which the importance of feature *combinations* is derived from the classification algorithm itself. A successful classification indicates the presence of important discriminating features and finding these features yields the required variable combination

The muscles playing important roles in tuning for different pointing directions are listed in Table 2C. They are all the deltoid muscles in the downward directions, along with the trapezius muscle. In the upward direction, the posterior deltoid no longer plays a key role, while two additional trunk muscles, the pectoralis major and latissimus dorsi contribute to tuning towards 0°, compared to 90°. This is in keeping with previous studies which have shown these muscles to have different activities as a function of pointing directions (Flanders 1992; Flanders et al, 1994, 196; Mira et al, 2021). The novelty in the current study is that we have used Machine Learning to pick out the muscles among the collection of recorded muscles which are most pertinent to altering pointing directions and highlighted when these changes occur. This is not a trivial consideration as this would increase the ease of visualization and model construction especially for more complex movements. This would then improve the characterization of movements in healthy subjects and hence draw attention to compensatory movement patterns that might indicate early onset of neuromuscular deficiencies. Another novel aspect of the current study was the introduction of the *LDA_distance_* variable that provided a means of understanding certain global characteristics of EMG alterations for pointing directions. For example, overall EMG adjustments for pointing in the upward direction were lower than those for the downward direction when compared to horizontal pointing. The higher *LDA_distance_* at 180° in Figure 3B demonstrates this. This may be due to the fact that, although horizontal and upward movements follow a classical tri-phasic burst organization (Hallett et al. 1975; Virji-babul et al. 1994), downward movements do not (Gaveau et al. 2021). Gaveau et al (2021) have provided several arguments to show that this is perhaps because gravity replaces the agonist burst to accelerate the arm downwards, thereby saving muscle effort. Downward movements therefore exhibit activation patterns that strongly deviate from the classical tri-phasic burst pattern (Gaveau et al. 2021; Poirier et al. 2022). Thus, the temporal organizations of upward and horizontal movements are more similar to each other than are those of downward and horizontal movements.

Concerning the roles of the individual muscles as seen in table 2C, the importance of the deltoid muscles in classification would be due to their their role in setting the direction for the different angles of pointing. It is striking that the posterior deltoid while playing a key role in tuning for downward pointing is less important in tuning for upward pointing. During the acceleration part of upward and horizontal (rightwards) movements, substantial activity of the posterior deltoid muscle is needed to respectively stabilize the joint and accelerate the arm (Gaveau et al. 2021). During downward movements, however, this muscle remains largely silent. Again, the fact that the effort from the posterior deltoid is replaced by gravity torque, during a downward movement, makes the pattern of this muscle very different from its activation during a horizontal one. Regarding why the latissimus dorsi is useful in classifying upward but not downward movements compared to horizontal ones, this may be due to the fact that this muscle is engaged in stabilizing the shoulder joint and decelerating the arm when moving upwards. On the contrary, this muscle is less if any activated during downward (its effort is replaced by gravity torque) and horizontal movements (it is perpendicular to the plane of motion). The relatively low prominence of muscles such as the biceps and triceps acting around the elbow joint might be due to the protocol of the experiment in which elbow rotation was discouraged. This is in contrast to the experiments conducted by the Flanders group (Buneo et al, 1994, 2008; Flanders et al, 1991, 1992, 1996) where a pointing protocol with elbow rotations was involved. The important role played by the trapezius muscle for modulation in both directions is in agreement with reports by other groups that along with the anterior deltoid, it plays an important role in shoulder orientation for pointing direction (Tokuda et al, 2016; Sabatini et al, 2002). It should be noted that none of the differences in contribution to classification here is due to differences in EMG amplitude as this variable is normalized (see Methods).

In conclusion we will say that in the era of big data, Machine Learning Classification with LDA appears to be a useful tool which can complement currently available techniques like univariate statistics and PCAs in the study of motor control. Univariate statistics which are the most widely employed analytical tool in motor control studies have been extremely useful in confirming or rejecting pre-conceived notions on important variables. This technique is less viable in the face of big volumes of data and less open to the possibility of previously unexpected influences. The use of ensemble techniques like PCA, used to find synergies have been extremely useful in tackling the problem of the number of degrees of freedom in the motor system (d’Avella et al, 2010, 2011; Delis et al, 2018). They have a completely different goal from the current paper. They aim to construct a common framework from which to describe various types of movement while the goal of the current project is to find differences. It should also be pointed out that the two methods are not incompatible. Once synergies are constructed, Machine Learning classification can then be used to assign classes based on the synergies.

## Acknowledgments

This research has been funded in part by the Institut National de la Santé et de la Recherche Médicale (INSERM) of France and by the Conseil Régional de Bourgogne.

## Notes

### Competing Interest Statement

The authors have declared no competing interest.

